# Neutrophil TLR2 signaling promotes lipid accumulation and vascular plaque growth

**DOI:** 10.1101/2025.07.09.663961

**Authors:** Anudari Letian, Kyrlia Young, Anna Pi, Michelangelo Gonzatti, Regan Volk, Isha Sharma, Clayton Baker, Bérénice A. Benayoun, Balyn W. Zaro, Amber N. Stratman, Emily L. Goldberg

**Affiliations:** Department of Physiology, University of California, San Francisco, San Francisco, CA, USA; Department of Pharmaceutical Chemistry, The Cardiovascular Research Institute, University of California, San Francisco, CA, USA; Leonard Davis School of Gerontology, University of Southern California, Los Angeles, CA 90089, USA; Molecular and Computational Biology Department, USC Dornsife College of Letters, Arts and Sciences, Los Angeles, CA 90089, USA; Cancer Biology Department, USC Keck School of Medicine, Los Angeles, CA 90033, USA; Pharmacology and Pharmaceutical Sciences Department, USC Mann School of Pharmacy, Los Angeles, CA 90033, USA; Department of Cell Biology and Physiology, Washington University in St. Louis School of Medicine, St. Louis, MO, USA; Chan-Zuckerberg Biohub, San Francisco, CA, USA

## Abstract

Neutrophils are short-lived cells that are produced by the billions every day to circulate throughout the body and surveil all tissues. They are a key component of the innate immune system that play essential roles in antimicrobial immunity, but can also instigate sterile inflammatory diseases like gout and cancer. Immunometabolic paradigms that were developed by studying T cells and macrophages establish that cellular metabolic programming dictates immune function. Neutrophils have long been known as glucose-reliant and highly glycolytic. But surprising neutrophil heterogeneity has recently been described, and roles for lipids have been reported in both granulopoiesis and mature neutrophil effector function. Therefore, we set out to uncover how neutrophils acquire lipids from their environment and how this influences their functionality in the context of lipotoxicity. We found that neutrophils take up both free fatty acids and complex lipoproteins, but that their uptake is regulated through different signaling pathways. Neutrophil lipoprotein uptake is inducible by certain TLR2 signals, and this causes neutrophils to depolymerize their actin fibers and stop moving. Using a mouse model of atherosclerosis, we show that neutrophils in the plaque are lipid-laden and that neutrophil-deficient mice are protected from atherosclerotic plaque growth. Lipoprotein uptake causes neutrophils to recruit macrophages, conditional ablation of TLR2 on neutrophils prevents their lipid uptake and storage, and these mice are also protected against atherosclerosis. Our work highlights an important understudied role for lipids in neutrophil biology, and the importance of studying different lipid classes and different signaling pathways in neutrophils as compared to other myeloid populations.

---

Neutrophils are short-lived innate immune cells that circulate throughout the body and can be recruited to any organ upon infection or injury (*1-3*). While best studied for their important role in host defense against bacterial and fungal pathogens, a mounting body of evidence also establishes roles for neutrophils in non-infectious chronic inflammatory diseases like metabolic disease and gout (*4-6*). Recently, neutrophils were reported to infiltrate adipose tissue as active participants in β3 adrenergic receptor-induced lipolysis (*7*) and cold-induced thermogenesis. All of these data point to neutrophils as key players in organismal physiology, homeostasis, and health.

Decades ago, neutrophils were shown to be highly glycolytic (*8, 9*). As short-lived cells, this dogma has persisted and remained the primary focus of studies on neutrophil metabolism. Indeed, the majority of mature neutrophil ATP production comes from glycolysis. Neutrophils even synthesize and store their own glycogen, which is believed to serve as a fuel source in hypoxic nutrient-replete tissues (*10-13*). But a growing body of literature shows that neutrophils are more metabolically flexible than previously assumed, and this supports their diverse effector functions, similar to the immunometabolic regulation that is well described in T cells and macrophages. Lipids also play an important role in neutrophil biology (*14*). Neutrophils can store lipid droplets and access free fatty acids from these stores, either through autophagy or lipolysis, has been linked to successful granulopoiesis and tumor metastasis (*15, 16*). Likewise, blocking neutrophil lipid uptake through the FATP2 transporter slows tumor growth (*17*). Depleting cellular cholesterol or treating neutrophils with statins increases their NETosis (*18-20*). Beyond lipid storage, neutrophils also metabolize fatty acids and deletion of CPT1a or pharmacological inhibition of CPT1a with etomoxir influences bacterial killing (*21*), cell movement (*20*), and modestly reduces NETosis in combination with other metabolic substrates (*22*). In totality, this work establishes that metabolic and non-metabolic fates of lipids need to be carefully considered in regulation of neutrophil biology.

When we reanalyzed prior neutrophil transcriptomics datasets, we became intrigued by the high levels of triglyceride synthesis and lipid storage genes like diacylglycerol O-acyltransferase 1 (Dgat1) and perilipins (Plin2, Plin3) but very low expression of de novo lipogenesis genes like Fatty Acid Synthase (Fasn) and Acetyl-CoA carboxylase (Acaca) (*23, 24*). We also verified that public databases Immgen and ImmPRes (*25, 26*). While fatty acid synthesis is essential in granulopoiesis, most studies in mature neutrophils have focused on lipid utilization (*15, 22, 27*). Therefore, we initiated this study to explore the mechanisms of lipid acquisition in neutrophils and its functional consequences. We found that neutrophils take up a variety of lipids both *in vitro* and *in vivo*. Moreover, we found that lipid uptake can be stimulated in neutrophils downstream of TLR signaling, but this is lipid class-dependent. Our data also indicate that different receptors might be used to take up different lipids and that neutrophil lipid uptake is CD36-independent. Like what has been shown after pathogen phagocytosis in neutrophils and other myeloid cells (*28-31*), lipid-containing neutrophils also migrate less and our data indicate this is due to actin depolymerization. Finally, to establish the physiological relevance of neutrophil lipid uptake, we tested the role of neutrophils in atherosclerosis. Neutrophil-deficient mice were protected from developing atherosclerotic plaques, in agreement with prior work (*32, 33*), and mice with conditional TLR2 deficiency on neutrophils also had smaller plaques. Altogether, our study highlights an important and relevant role for neutrophil lipid uptake and regulation in sterile inflammatory diseases of lipotoxicity.

## Results

We reanalyzed two prior independent neutrophil transcriptomics datasets and found that neutrophils express high levels of the triglyceride synthesis gene *Dgat1*, and the lipid droplet proteins *Plin2* and *Plin3*, but exceptionally low levels of fatty acid synthesis genes *Fasn* and *Acaca* (**Fig 1A, S1A-D**) (*23, 24*). These data suggested to us that neutrophils might acquire lipids from the environment and then package them for storage. To compare lipid uptake in different white blood cell populations, we injected mice retro-orbitally with a fluorescent palmitate-BODIPY, as an abundant, model long-chain fatty acid that we could track using flow cytometry. To our surprise, we found that neutrophils readily took up the palmitate-BODIPY compared to T cells, B cells, and monocytes in the blood and spleen (**Figs 1B**). To further examine if neutrophils take up other classes of lipids, we used an *in vivo* acute peritonitis model combined with fluorescent lipid treatment to compare neutrophil lipid uptake to that of other recruited phagocytes. In these experiments, we injected zymosan into the peritoneal cavity to recruit neutrophils and other myeloid cells. Three hours later, we injected the same mice with fluorescent oxidized low-density lipoprotein (oxLDL)-DiI to use flow cytometry to measure uptake of this representative rare lipid species. Like we observed with palmitate, neutrophils robustly took up the oxLDL-DiI, even compared to other recruited macrophages and monocytes (**Figs 1C-E**). We used confocal microscopy to confirm the intracellular localization of both these fluorescent lipids (**Fig 1F**). Interestingly, we observed that oxLDL uptake, but not palmitate or even unmodified LDL uptake, led to lipid droplet formation in neutrophils (**Fig. S1E**), indicating that different lipids have different fates within neutrophils. Finally, we obtained slides of mouse atherosclerotic plaques from neutrophil fate-mapped mice (S100a8-Cre x Ai14.TdTomato, visualized in magenta) and confirmed that lipid-laden neutrophils were readily detected in atherosclerotic plaques (**Fig 1G**). Importantly, we used CD68 staining to verify that these were not macrophage foam cells (visualized in white), which are abundant and well-known lipid reservoirs in plaques. These data indicate that neutrophils can actively scavenge for lipids from their environment.

**Figure 1.**
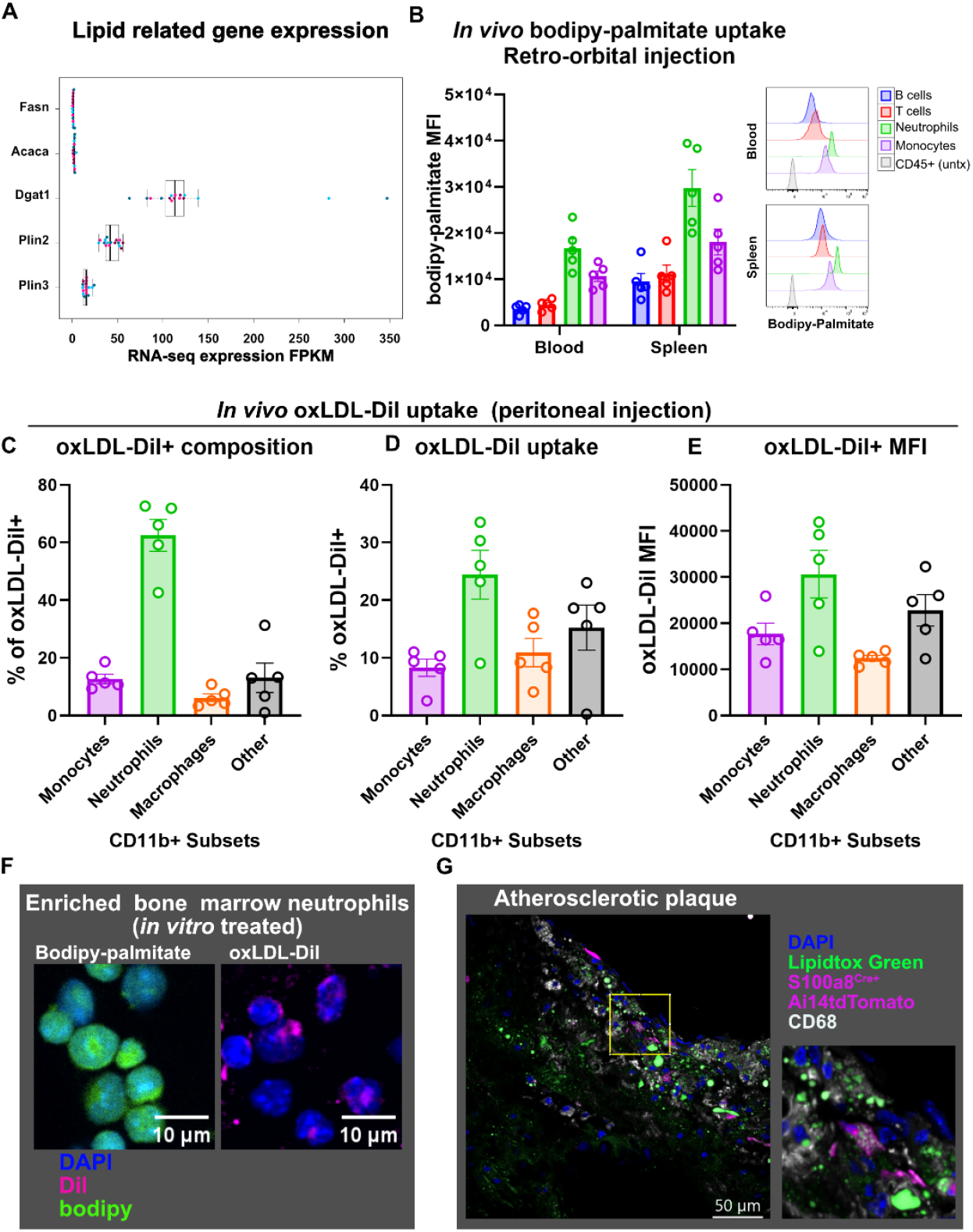
Neutrophils robustly take up a variety of lipids. (**A)** Bulk neutrophil RNA-Seq expression of fatty acid synthesis and lipid storage genes. Blue dots represent male mice, pink dots represent female mice. **(B**) *In vivo* uptake of intravenous injected bodipy-palmitate. Immune cells from the blood and spleen were analyzed via flow cytometry. Bodipy MFI was quantified and histogram of bodipy-intensity is plotted for the specified lineages. Populations were defined as B cells (CD45^+^, B220^+^), T cells (CD45^+^, CD3^+^), neutrophils (CD45^+^, CD11b^+^, Ly6C^med^, Ly6G^+^), and monocytes (CD45^+^, CD11b^+^, Ly6C^high^, Ly6G^-^). (**C-E**) *In vivo* oxLDL-Dil uptake in zymosan-elicited peritoneal cells. (**C**) Composition of oxLDL-Dil+ cells. (**D**) oxLDL-Dil uptake in specified populations. (**E**) oxLDL-Dil MFI of the oxLDL+ subpopulation within each cell type. Populations were defined as neutrophils (CD11b^+^, Ly6C^med^, Ly6G^+^), monocytes (CD11b^+^, Ly6C^high^, Ly6G^-^) and macrophages (CD11b^+^, F4/80^+^). (**F**) Representative confocal images of peritoneal cells following *in vitro* bodipy-palmitate and oxLDL-Dil uptake. (**G**) Atherosclerotic plaque section from S100a8Cre^+^ Ai14tdTomato mouse stained for CD68 and Lipidtox Green. For plots, each graph is a representative of at least three experiments where each data point is a replicate from one mouse. Data are expressed as mean ± SEM.

If neutrophil lipid uptake is regulated by immune signals. We started with receptor signaling on the plasma membrane because we were measuring extracellular lipid uptake and focused first on toll-like receptor (TLR) signaling. We used Pam3CSK4 and FSL-1 to activate TLR2, LPS to activate TLR4, and commercially sourced *Salmonella typhimurium* flagellin to activate TLR5. To simulate the effects of TLR3, TLR7, and TLR9 activation, we also treated cells with interferon β (IFN-β). We observed that palmitate-BODIPY uptake was decreased under all these conditions (**Fig 2A**). In contrast, uptake of oxLDL-DiI followed a different pattern. We found that neutrophil lipoprotein uptake could be induced by activating TLR2 and TLR5, but neither TLR4 nor IFN-β stimulated lipid uptake (**Fig 2B**). We were especially intrigued to find that different TLR2 agonists had different effects on lipoprotein uptake. Specifically, the TLR1/2 activator Pam3CSK4 did not alter oxLDL uptake, whereas the TLR2/6 activator FSL-1 significantly increased oxLDL uptake. These data indicate that lipid uptake in neutrophils is inducible, downstream of certain TLR pathways, and may be uniquely regulated for individual classes of lipids.

**Figure 2.**
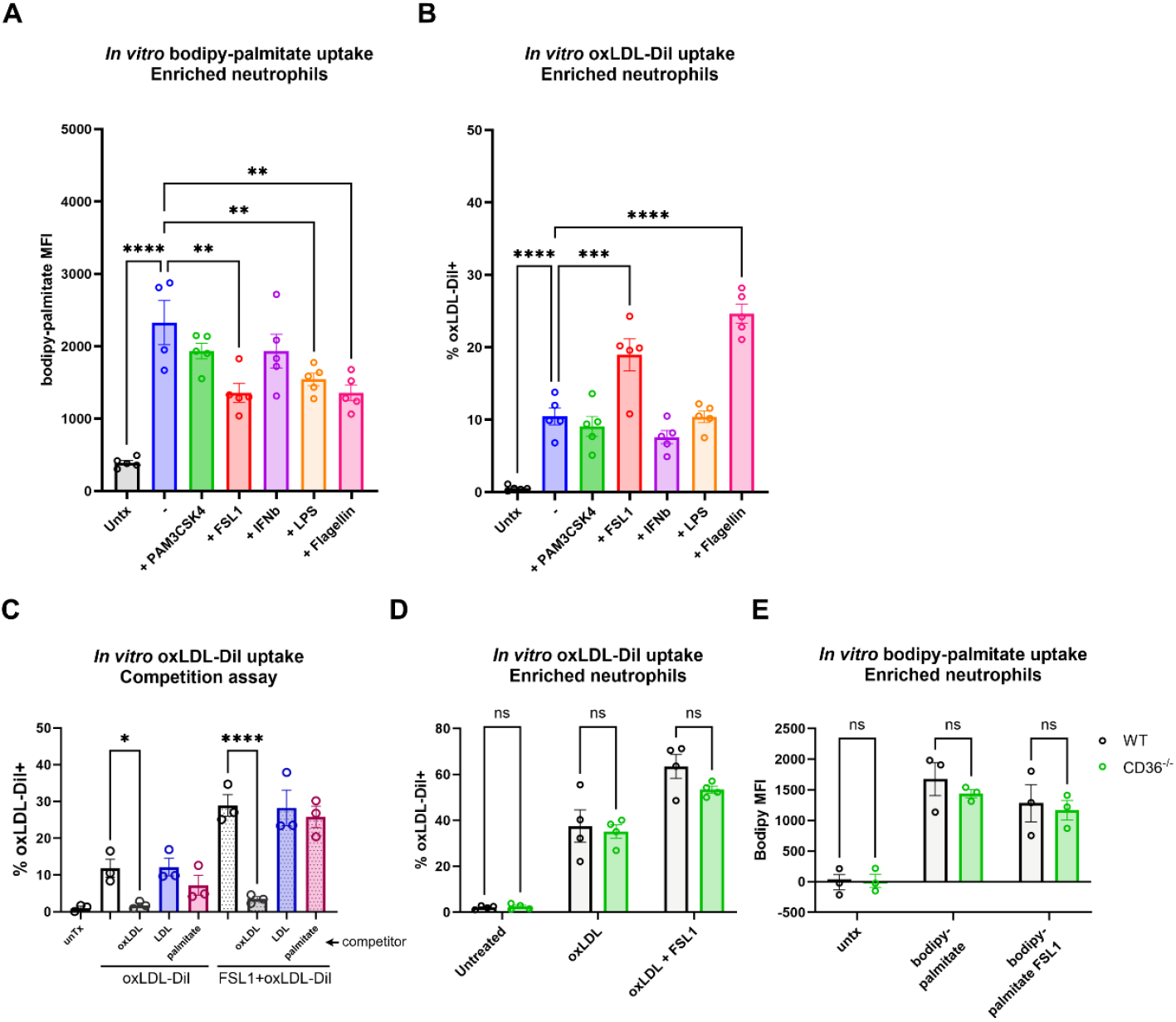
Neutrophil lipid uptake is induced by TLR signaling and is CD36-independent. (**A-B**) *In vitro* fluorescent lipid uptake under baseline and TLR-stimulated conditions. Bone marrow enriched neutrophils (CD45^+^, Ly6G^+^) were incubated with (**A**) bodipy-palmitate, (**B**) and (**B**) oxLDL-Dil simultaneously with the specified TLR ligands and analyzed via flow cytometry. (**C**) *In vitro* oxLDL-dil uptake competition assay using molar excess of non-fluorescent lipids. *In vitro* (**D**) oxLDL-Dil and (**E**) bodipy-palmitate uptake in neutrophils isolated from WT or CD36 knock-out mice. Each data point is a biological replicate experiment using neutrophils pooled from two mice. Data are expressed as mean ± SEM. For (**A-C**), statistical differences were calculated by one-way ANOVA. For (**D-E**), statistical differences were calculated by two-way ANOVA. * ≤ 0.05, ** ≤ 0.01, *** ≤ 0.001, **** ≤ 0.0001

We next investigated how neutrophils take up extracellular lipids. To test if lipid uptake is receptor-mediated, we developed a competition assay in which neutrophils were treated with oxLDL-DiI in the presence or absence of a molar excess of non-fluorescent lipid species. Non-fluorescent oxLDL blocked oxLDL-DiI uptake, but oxLDL-DiI uptake was unaffected by LDL or palmitate co-treatment (**Fig 2C**). These data indicate that oxLDL is likely taken up through a unique receptor, or at minimum, through a saturable mechanism. CD36 is a lipid scavenger receptor previously shown to mediate oxLDL uptake in macrophages and monocytes (*34-36*). However, neutrophils from *Cd36*^-/-^ mice also had normal oxLDL-DiI and palmitate-BODIPY uptake (**Figs 2D-E, S2**). Taken together, these data suggest that neutrophil lipid uptake is receptor-mediated, that receptors are specific for certain lipid species, but that neutrophils might use unique receptors for lipid uptake compared to what has been studied in other cell types.

To unbiasedly assess the impact of lipid uptake, we again used our peritonitis model to purify lipid-positive and lipid-negative neutrophils to compare by LC-MS/MS (**Fig 3A**). Reactome Pathway Enrichment and Gene Ontology analyses revealed that oxLDL-positive neutrophils were significantly enriched in RhoGTPase signaling pathways and actin filament organization (**Fig 3B, S3**). When we further examined the proteins that were increased or decreased in the lipid-positive neutrophils, we were struck by the relative depletion of F-actin binding protein Myosin heavy polypeptide 11 (Myh11), the F-actin capping protein CAPZA2, and multiple Rab GTPases that are known to interact with F-actin but also be involved in endocytic trafficking (**Fig 3C**). In contrast, oxLDL-positive neutrophils were enriched for ARHGDIB and ARHGDIA, which inhibit activation of Rho proteins required for actin rearrangement, and ARHGAP45, which catalyzes GTP hydrolysis of Rho GTPases. Notably, oxLDL-positive neutrophils were also enriched for CORO1A, which was recently reported to be involved in neutrophil F-actin disassembly (*37*). This pattern of changes in actin remodeling proteins indicated that neutrophils that recently took up lipid might have compromised or depolymerized actin. To test this possibility, we isolated peritoneal cells from our peritonitis model and stained them with phalloidin to visualize F-actin fibers. As predicted by our proteomics data, oxLDL-positive neutrophils showed a dearth of phalloidin staining as compared to lipoprotein-negative cells (**Figs 3D, S4**). Cytochalasin D treatment blocked oxLDL-DiI uptake (**Fig 3E**), confirming that oxLDL uptake is an active process and indicating that the loss of F-actin occurs as a consequence of lipid uptake. We reasoned that actin disorganization could lead to impaired migration, as has been shown after neutrophil particle phagocytosis (*28*). Indeed, transwell migration assays of purified oxLDL-positive and oxLDL-negative neutrophils confirmed that lipid-positive neutrophils migrate less both spontaneously and towards target cells (**Fig 3F**). These data highlight that lipid particle uptake by neutrophils can have remarkable functional consequences.

**Figure 3.**
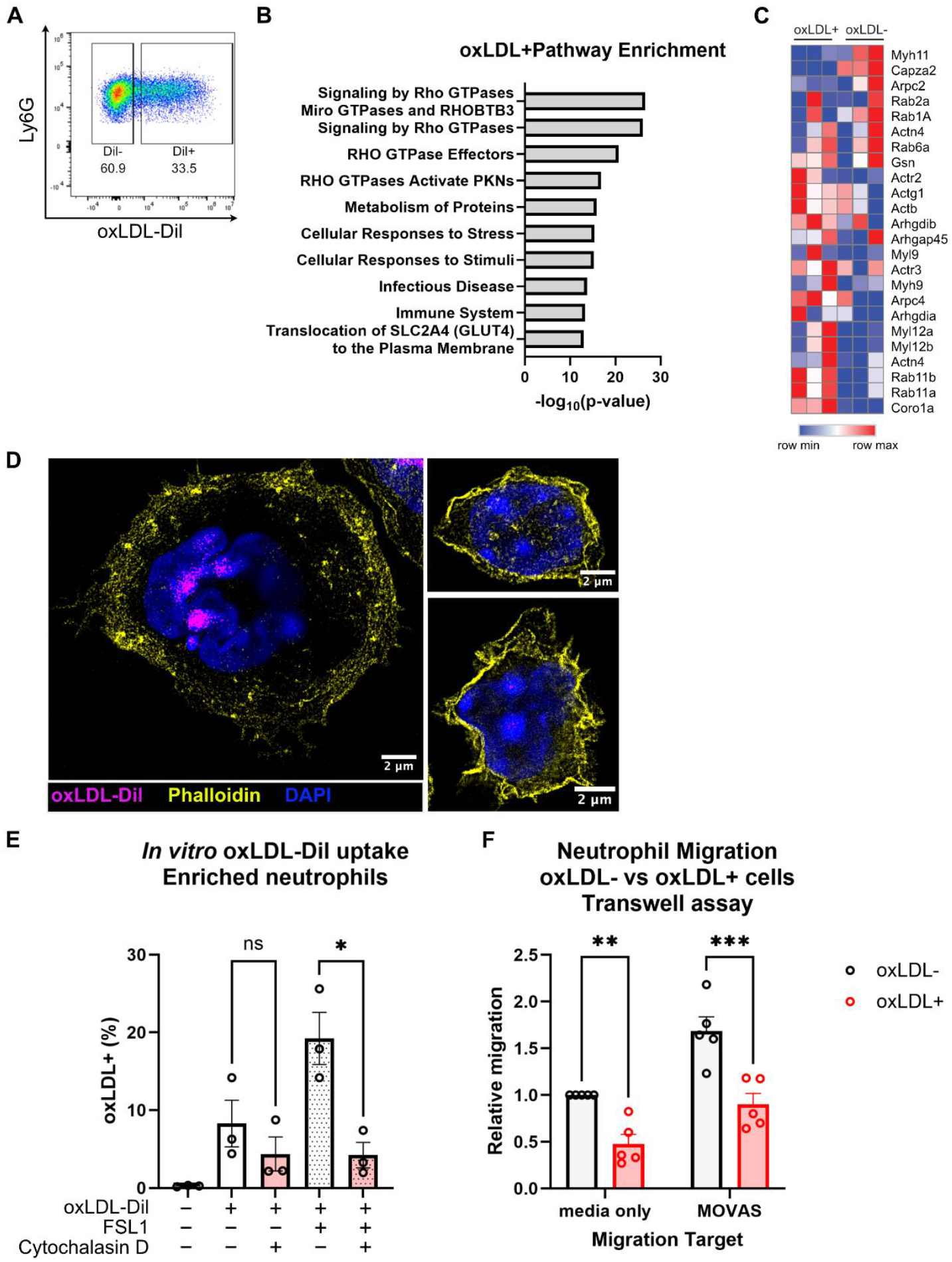
oxLDL uptake disrupts actin polymerization and causes neutrophil stalling. (A) Representative FACS plot of cell sorting strategy to isolate purified oxLDL^+^ and oxLDL^-^ neutrophils. (**B**) Reactome pathway analysis of proteins increased in oxLDL^+^ subset in the proteomics data. (**C**) Heatmap of selected differentially expressed proteins in oxLDL^-^ and oxLDL^+^ neutrophil populations. (**D**) 100x STED immunofluorescent imaging of peritoneal cells isolated from zymosan and oxLDL-Dil IP-injected mice and stained for phalloidin. Cells were also stained and checked for Ly6G expression, which is not shown for clarity of phalloidin stain. Representative oxLDL^+^ (left) and oxLDL^-^ (right) cells are shown. (**E**) *In vitro* oxLDL-Dil uptake in bone marrow enriched neutrophils treated with cytochalasin D. (**F**) Transwell migration assay of oxLDL^-^ and oxLDL^+^ neutrophils sorted from peritoneal cells following zymosan and oxLDL-Dil IP injection. (**E-F**) All symbols represent independent experiments. Data are expressed as mean ± SEM. Statistical differences were calculated by one-way ANOVA for F and two-way ANOVA for G. * ≤ 0.05, ** ≤ 0.01,***≤0.001

Next, we sought to establish the physiological relevance of TLR2-regulated lipid uptake in neutrophils. Given that we already observed lipid-laden neutrophils in atherosclerosis and oxLDL is an atherogenic lipid associated with foam cell formation, we returned to our inducible atherosclerosis model to test the significance of neutrophil-lipid interactions *in vivo*. We used an adeno-associated virus to specifically induce a gain-of-function mutation in hepatocyte PCSK9 (AAV8-mPCKS9-D377Y) (*38*). PCSK9 is responsible for low-density lipoprotein receptor (LDLR) recycling, and this mutation prevents LDLR recycling back to the plasma membrane, rendering hepatocytes functionally LDLR-deficient and driving hypercholesterolemia that is necessary for atherosclerotic disease and plaque formation. This model provided the added advantage that all lipid handling machinery in neutrophils would remain intact as compared to classical *Ldlr*^-/-^ or *ApoE*^-/-^ mouse models often used to study atherosclerosis (*39-42*). Using our neutrophil fate-mapping mice, we found that neutrophils are extremely rare in the heart and aorta in lean healthy mice, but began infiltrating these tissues within weeks after induction of atherosclerosis (**Fig S5A**). These experiments also revealed that only the neutrophils that had infiltrated the plaques became lipid-laden, and that neutrophils outside the plaque, within other regions of the heart, did not contain lipid droplets (**Fig S5B**). Importantly, although other immune cell types may express S100a8 in atherosclerosis, our fate mapping data confirm that at least 80% of the Cre+ cells in atherosclerotic hearts and aorta are neutrophils, and that no other CD11b+ myeloid lineages had S100a8-Cre activity (**Fig S5C**,**D**).

To confirm that neutrophils are important in atherosclerosis pathophysiology, we induced atherosclerosis in neutrophil-sufficient and neutrophil-deficient littermates (using S100a8-Cre+/-mice crossed to lox-stop-lox DTA mice). In agreement with prior work that used an antibody to acutely deplete neutrophils (*33*), we found that mice with genetic constitutive neutrophil depletion were protected from atherosclerosis (**Figs 4A-D, S6**). Neutrophil-deficient mice had smaller plaques in their aortic roots. We used CD68 and Acta2 staining to assess plaque macrophage and smooth muscle cells and found that while the plaques in neutrophil-deficient mice were smaller, their cellular composition was otherwise normal (**Figs 4E-H**). Of note, because we saw differences in body weights of the Cre+ mice that we were not expecting, we did confirm that the presence of S100a8-Cre itself does not affect plaque size in control cohorts (**Fig S7**). After confirming that TRL2-deficient neutrophils lack inducible oxLDL uptake by TLR2 agonists (**Fig 5A**), we crossed the S100a8-Cre mice to *Tlr2*^fl/fl^ mice to conditionally ablate TLR2 on neutrophils. We confirmed that isolated bone marrow neutrophils from these conditional knockout mice failed to form lipid droplets (**Fig 5B**), so we used them to further evaluate the role of neutrophil TLR2 signaling in atherosclerosis. Importantly, neutrophil-specific TLR2 cKO mice also had smaller plaques as compared to their Cre-negative littermates (**Figs 5C-F**). Previous work identified non-hematopoietic TLR2 expression as being more important in atherosclerosis (*43, 44*), but neither of these studies specifically tested neutrophils.

**Figure 4.**
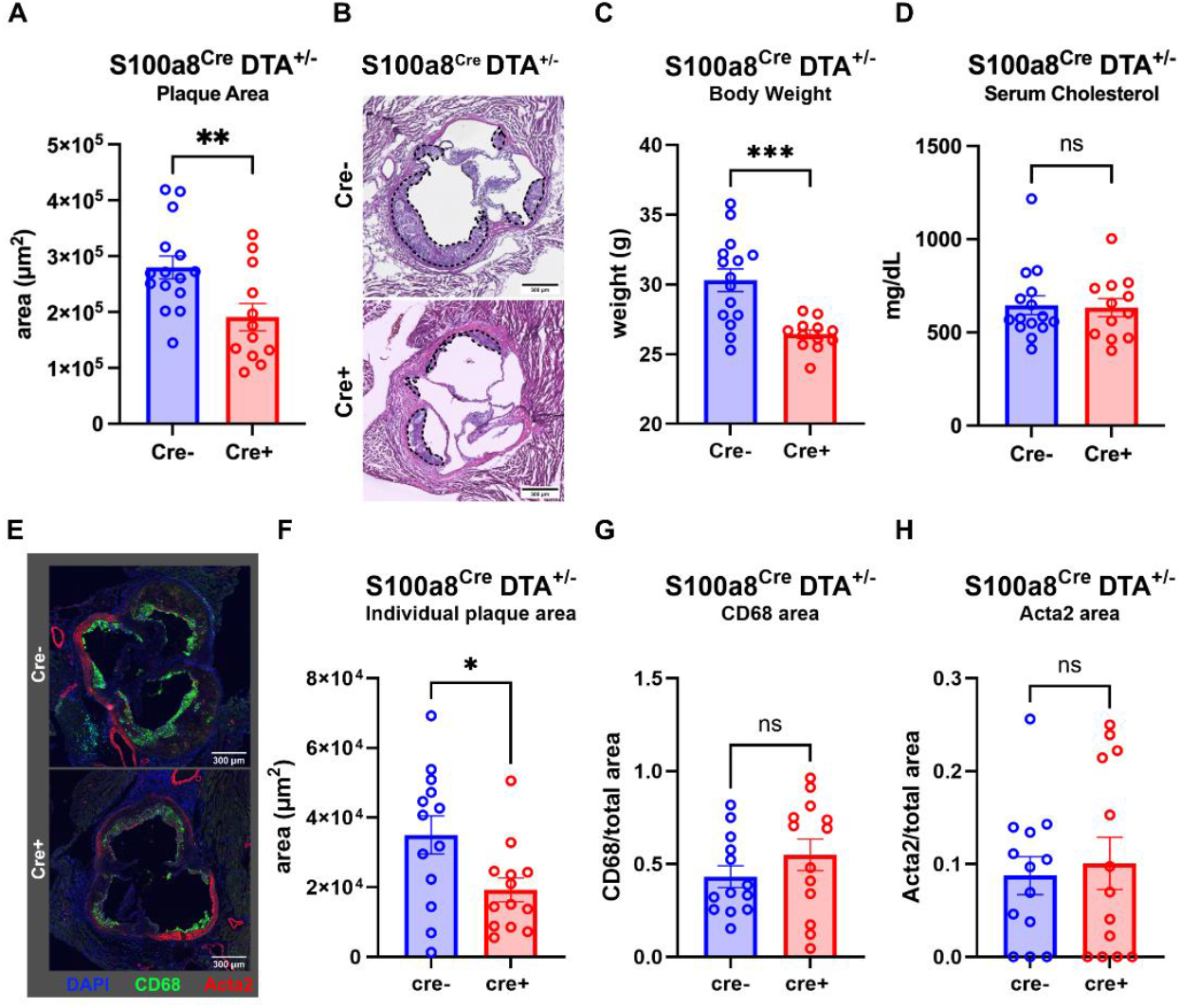
Neutrophil-deficient mice are protected from atherosclerosis. Atherosclerosis was induced in male mice via combination of AAV8-PCSK9 injection and high fat high cholesterol diet. **(A)** Plaque area quantification, (**B**) representative images of H&E stain of aortic root, and (**C**) mouse body weights at 12 weeks of disease are shown for S100a8^Cre+^ DTA^+/-^ and littermate control mice. (**D**) Serum cholesterol quantification at 4 weeks of disease for the respective genotypes. Quantified plaques are highlighted with dashed lines. (**A-D**) Data are compiled from three independent mouse cohorts, each data point on the graph represents a single animal. (**E**)Representative confocal images of aortic root section stained with CD68 and Acta2 from S100a8^Cre+^ DTA^+/-^ and littermate control mice at 12 weeks of atherosclerosis induction. Quantification of (**F**) plaque area, (**G**) CD68 area normalized to plaque area, (**H**) ACTA2 area normalized to plaque area for each individual plaque is shown. (**E-H**) Data are from one representative mouse cohort. Data are expressed as mean ± SEM. Statistical differences were calculated by t-test. * ≤ 0.05, ** ≤ 0.01, *** ≤ 0.001

**Figure 5.**
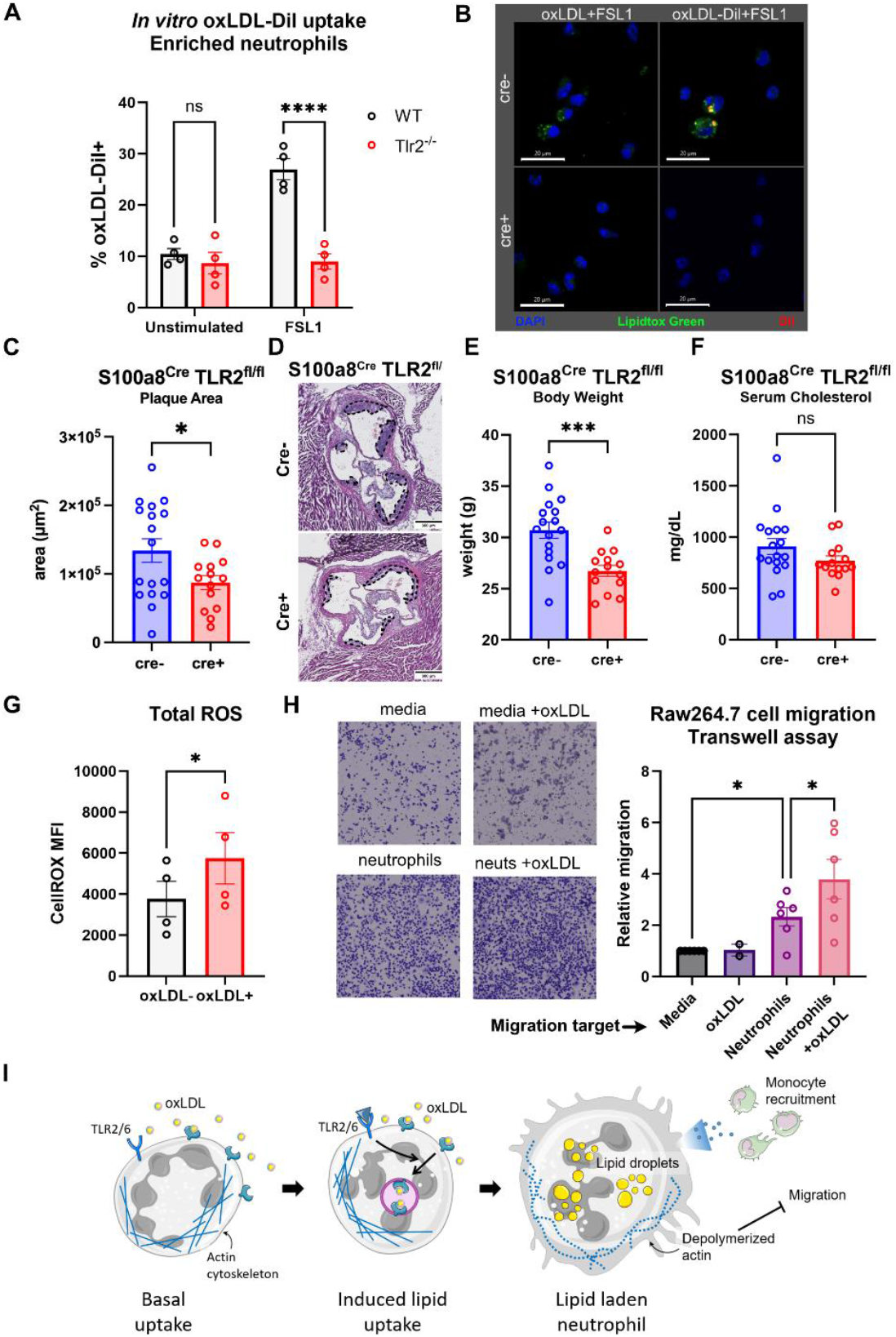
TLR2 ablation in neutrophils protects against atherosclerosis. (**A**) *In vitro* oxLDL-Dil uptake in neutrophils isolated from WT or TLR2 knock-out mice. Each data point is a replicate experiment. (**B**) Lipidtox Green stain in neutrophils, isolated from S100a8Cre^+^ TLR2^fl/fl^ and littermate control mice that were treated overnight with oxLDL or oxLDL-Dil. (**C-F**) Atherosclerosis characterization in S100a8Cre^+^ TLR2^fl/fl^ and littermate control mice. (**C**) Plaque area quantification, (**D**) representative images of H&E stain of aortic root. Quantified plaque regions are highlighted with dashed lines. (**E**) Mouse body weights at 12 weeks post-atherosclerosis induction. (**F**) Serum cholesterol quantification at 4 weeks of disease. Data is compiled from two independent mouse cohorts, and each data point represents a single animal. Data are expressed as mean ± SEM. (**G**) CellRox in oxLDL^-^ and oxLDL^+^ neutrophils isolated following zymosan and oxLDL-Dil IP-injection were analyzed by flow cytometry. Each symbol represents an independent animal and oxLDL^+^ and oxLDL^-^ CellROX MFI was compared within the same animal by paired t-test. (**H**) RAW264.7 cell transwell migration towards media and plated neutrophils with or without oxLDL. Each symbol represents an independent experiment. (**I**) Summary model figure for proposed mechanism and consequences of neutrophil lipid uptake. For (**A**), statistical differences were calculated by two-way ANOVA. For (**C, E, F, G**) statistical differences were calculated by t-test. For (**H**) statistical differences were calculated by one-way ANOVA. * ≤ 0.05, *** ≤ 0.001, **** ≤ 0.0001 aorta in lean healthy mice, but began infiltrating

Taken together, our data suggest that differences in signaling pathways across different cell types may be an important aspect of atherosclerosis that needs further exploration. To understand possible mechanisms of how lipid uptake in neutrophils might contribute to atherosclerotic disease, we assessed ROS production, a common feature of atherosclerosis and a well-known effector function of neutrophils. We compared total ROS content between oxLDL-positive and oxLDL-negative neutrophils from our *in vivo* peritonitis model and found that oxLDL-positive neutrophils had higher ROS content (**Fig 5G**). oxLDL treatment also induced neutrophils to recruit more macrophages in a transwell migration assay (**Fig 5H**). These data show that neutrophils could perpetuate plaque growth throughout disease by chronically encountering and acquiring tissue lipids, stalling their movement, and recruiting additional disease-promoting immune cells to the lesions (**Fig 5I**).

## Discussion

Our data reveal an interesting paradigm in neutrophils that lipid acquisition can be activated by immune pathways like TLR signaling and that different lipids are under unique regulatory mechanisms. The specificity of TLR2/6 heterodimer activators like FSL1 and zymosan for inducing lipid uptake, as compared to TLR1/2 activator Pam3CSK4, demonstrates that different downstream signaling pathways from TLR2 may tune responses to different stimuli. CD36 has been described as a TLR2-inducible oxLDL receptor, but our data show that neutrophil lipid uptake does not require CD36, implying that additional lipid co-receptors are present.

At this time, it is unclear why lipid uptake leads to loss of F-actin in neutrophils. Prior work has shown that neutrophils stop migrating after larger particle phagocytosis, but had normal F-actin staining, and this was instead due to changes in biophysical membrane properties (*28*). Neutrophils reportedly have phagocytic preference for certain target shapes and sizes (*45*) and we wonder if the physical properties of different particles play a role in downstream neutrophil consequences. We propose that different lipid classes could have a similar paradigm. While our data do not suggest that lipids are phagocytosed, we speculate that altered migration may be a common downstream consequence of particle engulfment by neutrophils, irrespective of particle identity. We further hypothesize that neutrophils may have multiple pathways to stop migrating upon cargo uptake, and that these pathways could change depending on particle size, particle identity, or perhaps the route of particle entry.

In the future, it will be important to determine the fates of different lipids in neutrophils and how this might be influenced by different inflammatory conditions. ^13^C-palmitate tracing has shown that fatty acids are used metabolically and contribute to TCA cycle intermediates (*20*). However, mitochondrial metabolism and specifically fatty acid oxidation are estimated to account for less than 10% of ATP production in neutrophils (*10*). One possible fate for lipids could be bioactive lipid production like prostaglandins, eicosanoids, or leukotrienes. Alternatively, lipids could be stored in lipid droplets, although the purpose of lipid droplets in mature neutrophils remains elusive. Given our data that the uptake of different classes of lipids is uniquely regulated, how different classes of lipids might feed into different pathways, or be involved in different immune responses, should also be explored in future studies. Many of the paradigms in metabolic control of immune function were established in T cells and macrophages. Our data, combined with prior work from others, show that neutrophils have more unique metabolic programming and nutrient sensing than previously realized, and should be more carefully studied for how this intersects with their diverse effector functions.

There is a lot of correlative evidence that neutrophils are associated with cardiovascular disease, and multiple studies have shown evidence of neutrophils in human and mouse atherosclerotic plaques (*46-54*). Previously, neutrophil depletion in early but not late-stage disease in atherogenic *ApoE*^-/-^ mice showed slowed plaque growth (*33*). Our data show that depleting neutrophils throughout the disease also protects against plaque growth. We are especially intrigued by our data that neutrophils in the plaque, but not neutrophils more distal in the heart parenchyma, contained lipid (**Fig S5B**). This raises the question whether a local endogenous TLR2 signal may stimulate lipid uptake in the plaque. Notably, this possibility was previously raised by Mullick et al., in 2005 (*44*). We must note, however, that our data do not directly implicate neutrophil lipid uptake in atherosclerosis, and this may be difficult to resolve until the relevant receptors are identified. The overall plaque composition in neutrophil-deficient mice is similar to littermate control neutrophil-sufficient mice. This, combined with our data that neutrophils continue to infiltrate the aorta with increasing numbers throughout the course of disease, suggests that they are continuously recruited to disease regions and play a pivotal role in perpetuating plaque development. Of note, the TLR2 cKO phenotype is not as strong as the complete neutrophil deficiency, indicating additional pathways may be involved. This possibility is supported by our lipid uptake in Figure 2, which shows that TLR5 also induces lipoprotein uptake, a possibility that can be followed up on in future experiments. It is also possible that additional unknown signals can stimulate lipid uptake in neutrophils. Altogether, our data point towards a model in which neutrophils are recruited to the atherogenic vascular wall where they encounter and take up lipid that causes them to stop moving, they then recruit other disease-promoting cells like macrophages, and possibly contribute directly to disease pathophysiology. How neutrophils participate in other chronic sterile inflammatory diseases of lipotoxicity, and how this intersects with their antimicrobial functions upon infection, also remain important outstanding questions that need more attention.

## Materials and methods

### Animals

All procedures were approved by the UCSF Institutional Animal Care and Use Committee. All mice were housed under specific pathogen-free conditions on a 12-hour light/dark cycle. Neutrophil-deficient mice were bred and housed in an SPF room that was additionally negative for murine norovirus and helicobacter. All mouse strains were on a C57BL/6J background and obtained from The Jackson Laboratory. CD36^-/-^ (JAX #019006), TLR2^-/-^ (JAX #004650) were obtained from The Jackson Laboratory and bred in house for experiments. To produce neutrophil-depleted mice, S100a8-Cre^+/-^ (JAX #021614) mice were crossed with DTA^lsl^ (JAX #009669) mice, yielding S100a8-Cre^+^ DTA^+/-^ (neutrophil-depleted) and S100a8-Cre^-^ DTA^+/-^ (littermate controls). To generate neutrophil-specific TLR2 knockout mice, S100a8-Cre^+/-^ mice were crossed with TLR2^fl/fl^ mice (JAX #037092), resulting in S100a8-Cre^+^ TLR2^fl/fl^ (neutrophil-specific TLR2 KO) and S100a8-Cre^-^ TLR2^fl/fl^ (controls). Additionally, S100a8-Cre^+/-^mice were crossed with tdTomato^lsl^ mice (JAX #007914) to obtain animals with red fluorescently labeled neutrophils. Female mice were used for lipid uptake experiments, while male mice were used in the atherosclerosis model.

### *In Vivo* Lipid Uptake

Female C57BL/6J mice aged 8–12 weeks were used for *in vivo* lipid uptake assays. For peritoneal lipid uptake experiments, mice received an intraperitoneal (i.p.) injection of either zymosan (100 µg in 200 µl PBS). Three hours later, the specified fluorescently labeled oxLDL-Dil was injected i.p. on the contralateral side (5 µg in 100 ul PBS). Two hours following lipid injection, mice were euthanized, and peritoneal cells were collected by lavage using 5 ml of ice-cold PBS. Cells were then washed and prepared for flow cytometric analysis. For intravenous lipid uptake assays, BODIPY-palmitate was administered via retro-orbital injection (0.01 µmol in 100 ul PBS). Twenty minutes post-injection, mice were euthanized, and blood was collected via cardiac puncture. Spleens were harvested into RPMI supplemented with 5% FBS, mechanically dissociated, and processed for flow cytometry.

### *In Vitro* Lipid Uptake

Neutrophils were isolated from the femurs and tibias of 8–12-week-old female C57BL/6J WT, TLR2^-/-^ and CD36^-/-^ mice using the StemCell Technologies magnetic negative selection kit (Catalog #19762), following the manufacturer’s protocol. One million cells were plated per condition in serum-free Opti-MEM media. Cells were treated with the respective lipid probes (oxLDL-Dil 5µg/mL, bodipy-palmitate 1µM) and TLR agonists for 3 hours (PAM3CK9 5ng/mL, FSL1 10 ng/mL, IFNb 0.1 ng/mL, LPS 0.1µg/mL, Flagellin 300 ng/mL), except for bodipy-palmitate, which was added in the final 30 minutes of the 3-hour incubation. Cytochalasin D was used at 20µM. After incubation, cells were scraped, washed twice with ice-cold PBS, and processed for flow cytometric analysis. For lipid uptake competition assay, neutrophils were isolated as above and co-incubated with fluorescent oxLDL-Dil (4 µg/mL) and a molar excess of non-fluorescent lipid (12.5 µg/mL for LDL or oxLDL, and 5 µM for palmitate). For imaging, 1 million cells were plated onto chamber slides, allowed to adhere for 30 minutes, and then fixed in 4% PFA for 15 minutes. For lipid droplet formation experiments, bone marrow enriched neutrophils were plated with the respective lipid at 16 hours and fixed. Following fixation, cells were processed for subsequent imaging and staining.

### Atherosclerosis Model

For atherosclerosis experiments, 12-week-old male mice were used because they are more susceptible to hypercholesterolemia in the AAV8-PCSK9 model (*55*). Disease was induced by retro-orbital injection of 1×10^11^ or 2×10^11^ viral vector genomes of AAV8-mPCSK9-D377Y (Addgene #58376, virus made by Penn Vector Core) (*38*). A lower dose was used for two out of three cohorts of S100a8-Cre+ DTA+/-; subsequently, atherosclerosis cohorts were injected with the higher dose of virus. Following injection, mice were switched to a high-fat diet (Clinton/Cybulski rodent diet containing 60 kcal% fat and 1.25% added cholesterol). At 4 weeks post-atherosclerosis induction, serum was collected from a tail bleed for cholesterol quantification using the Cholesterol Quantitation Kit (Sigma, MAK043-1KT) according to the manufacturer’s protocol. Mice with serum cholesterol levels below 400 mg/dL at 4 weeks were considered metabolic non-responders and excluded from further analysis. (11 mice from first two cohorts of S100a8-Cre^+^ DTA^+/-^ mice were excluded. No subsequent cohorts that were injected with the higher dose of virus resulted in any mice with serum cholesterol levels below 400 mg/dL). For plaque characterization, mice were euthanized at the indicated time points and perfused with PBS. Hearts were harvested into RPMI supplemented with 5% FBS, fixed in 4% PFA at 4°C for 24 hours, then transferred to a 20% sucrose solution for 24 hours. After embedding in OCT, hearts were sectioned transversely at 7 µm thickness and aorta was sectioned at 10 µm to obtain cross-sections of the aortic root for further analysis.

### Microscopy

Aortic root sections were stained either with hematoxylin and eosin (H&E) (Abcam, Catalog #ab245880) or with antibodies for immunofluorescence. H&E-stained sections were imaged using a 4x objective on a Keyence microscope. Plaque area was quantified manually using ImageJ. For immunofluorescent staining, primary antibodies were applied at a concentration of 1:200, and secondary antibodies at 1:500, in PBS containing 5% fetal bovine serum (FBS) and 0.25 mg/mL saponin. Slides were incubated for 30 minutes at 37°C. LipidTox Green (1:200 for tissue sections, 1:400 for cells) was used to stain neutral lipids for 30 minutes at room temperature. For phalloidin stain, cells were permeabilized with 0.01% Triton for 5 minutes, washed with PBS, and stained with Phalloidin STAR 635 at 1:400 in PBS + 1% FBS for 30 min at room temperature. DAPI stain (0.5 µg/mL) was used for all microscopy analyses. Slides were imaged using a Zeiss spinning disk and a BC43 spinning disk confocal microscopes, or using Keyence BZ-X810 for widefield microscopy. For super resolution imaging, slides were imaged using a four detector Abberior STEDYCON STED unit. Images were acquired as single plane to obtain maximal X,Y resolution (∼30nm), using a Leica HC PL APO 100x/140 oil CS2 objective. STED lasers were set between 10-20% power, based on the sample. The antibodies used in these assays included CD68 (Abcam, ab125212) and ACTA2-Cy3 (Sigma Aldrich C6198).

### Flow Cytometry

Red blood cells (RBCs) were lysed using ACK buffer. Spleens were harvested, mechanically dissociated through a 70 µm sieve, and RBCs were again lysed with ACK buffer. The resulting single-cell suspensions were washed twice with ice-cold PBS. For heart and aorta flow cytometry, mice were euthanized and perfused with 10 mL of PBS. Heart and aorta were dissected and stored in RPMI + 5% FBS on ice. For heart, tissue surrounding the aortic root was resected, minced finely. Both heart and aorta were digested in a shaking waterbath at 37°C 250 rpm for 50 minutes. Digestion cocktail for aorta consisted of 450 U/ml collagenase type I, 125 U/ml collagenase type XI, 60 U/ml hyaluronidase and 60 U/ml DNase1, Dulbecco’s phosphate buffered saline (DPBS) containing calcium. For heart, same cocktail was used except collagenase type XI was omitted. After digestion, cells were strained through 70 µm sieve, and single cell suspensions were washed twice with ice-cold PBS. For cells from digested tissues only, Zombie Aqua viability stain was used for 20 minutes on ice at 1:4000. Following PBS washes, cells were incubated with Fc block, followed by staining with an antibody cocktail for 20 minutes on ice. For ROS stain, CellROX Deep Red (Invitrogen) was incubated with live cells at 1:400 at 37°C for 30 minutes prior to antibody stain. After staining, cells were washed twice in PBS and acquired on Cytek Aurora or sorted on BD FACSAria™ Fusion. Data were analyzed on FlowJo™ v10.

### Transwell migration assay

For neutrophil transwell migration assay, zymosan elicited and oxLDL-Dil injected peritoneal neutrophils were sorted into oxLDL-positive and oxLDL-negative populations. 100k cells were plated in the top chamber of 3-micron pore transwell in 24-well plate in DMEM + 10% FBS. The bottom chamber contained either media only, or MOVAS cells (murine aortic smooth muscle cell line) plated at 15k per well the night before. Cell were allowed to migrate for 3 hours, and media from the bottom chamber was used to count migrated neutrophils in suspension. RAW264.7 cells were cultured in DMEM + 10% FBS. For migration, 50k RAW264.7 cells were plated in the top chamber in an 8-micron pore transwell plate in RPMI+ 10% FBS media. Either media alone, or 1 million bone marrow enriched neutrophils with or without oxLDL(10 µg/mL) were plated in the bottom well and cells were allowed to migrate for 24 hours. Wells were fixed with 4% PFA at room temperature, and stained in Crystal Violet for 20 min. Unmigrated cells were scraped off with damp cotton swab and membrane was cut out and mounted on slides. Slides were imaged on Keyence and analyzed with ImageJ. All cells were cultured in the indicated media supplemented with 1% antibiotic-antimycotic (Gibco, 15240-062) at 37 °C and 5% CO_2._

### Mass Spectrometry Sample Preparation

Zymosan-elicited neutrophils from our *in vivo* peritonitis experiment were sorted to obtain purified oxLDL-positive and oxLDL-negative Ly6G^+^ cells. Each sample was pooled from n=2 mice, and a total of 3 samples were analyzed. 125,000 cells were sorted from each population for each sample. Snap frozen cell pellets were allowed to thaw on ice before preparation with PreOmics iST96 kit. PreOmics iST lysis buffer was combined with boiling to solubilize membrane and soluble proteins.

### Mass Spectrometry Data Acquisition and Search Parameters

A nanoElute was attached in line to a timsTOF Pro equipped with a CaptiveSpray Source (Bruker, Hamburg, Germany). Chromatography was conducted at 40°C through a 25 cm reversed-phase C18 column (PepSep) at a constant flowrate of 0.5 µL min^-1^. Mobile phase A was 98/2/0.1% water/MeCN/formic acid (v/v/v), and phase B was MeCN with 0.1% formic acid (v/v). During a 108 min method, peptides were separated by a 3-step linear gradient (5% to 30% B over 90 min, 30% to 35% B over 10 min, 35% to 95% B over 4 min) followed by a 4 min isocratic flush at 95% for 4 min before washing and a return to low organic conditions. Experiments were run as data-dependent acquisitions with ion mobility activated in PASEF mode. MS and MS/MS spectra were collected with *m*/*z* 100 to 1700 and ions with *z* = +1 were excluded.

Raw data files were searched using PEAKS Online PEAKS Online Xpro 1.7.2022-08-03_160501 (Bioinformatics Solutions Inc., Waterloo, Ontario, Canada). The precursor mass error tolerance and fragment mass error tolerance were set to 20 ppm and 0.03 respectively. The trypsin digest mode was set to semi-specific and missed cleavages was set to 2. The mouse Swiss-Prot and TrEMBL database (downloaded from UniProt) totaling 86,618 entries was used. Carbamidomethylation was selected as a fixed modification. Deamidation (NQ) and Oxidation (M) were selected as variable modifications.

Raw data files and searched datasets are available on the Mass Spectrometry Interactive Virtual Environment (MassIVE), a full member of the Proteome Xchange consortium under the identifier: MSV000098121. The complete searched datasets are also available in Supplementary Table 1. To identify proteins of interest, we required at least 3 peptides to have been recovered from a particular protein, two of those proteins had to be unique, and the protein had to have been recovered in all three of the pooled mouse samples (irrespective of which neutrophil subset it was found in). Proteins that were then changed in the same direction in all three samples were further analyzed by Reactome Pathway Analysis (*56*) or Gene Ontology Analysis (*57-59*).

### Reanalysis of published neutrophil transcriptomic datasets

For bulk RNA-seq derived-analyses, we obtained processed gene-count matrices from bulk RNA-seq of purified mouse bone marrow neutrophils from Lu et al. 2021 (*23*). We then used DESeq2 v1.44.0 to calculate FPKM (fragments per kilobase of transcript per million mapped reads) values, enabling comparison of expression levels of multiple genes involved in triglyceride synthesis, lipid storage, and de novo lipogenesis. Briefly, count matrices were imported and cleaned by removing non-expression columns and standardizing gene and sample names. Sample metadata for age and sex was assigned based on experimental groups (young and old, male and female WT mice). Gene lengths were extracted and incorporated into the dataset for FPKM normalization. Expression data for selected lipid metabolism genes were then extracted for further analysis and visualization.

In parallel, to assess gene expression along bone marrow neutrophil maturation trajectories, we obtained a processed R data object containing pseudobulked single-cell RNA-seq (scRNA-seq) data of mouse bone marrow neutrophils from Kim et al. 2022 (*24*), generated using the muscat v1.5.2 framework. Pseudobulking was performed for each detected neutrophil subpopulations defined by Xie et al. 2020 (G2, G3, G4, G5a, G5b, G5c) (*60*). Pseudobulked counts were processed using DESeq2 v1.44.0 to compare variance-stabilized transformed (VST) counts, assessing the impact of neutrophil maturation on the expression of genes involved in triglyceride synthesis, lipid storage, and de novo lipogenesis. For each subpopulation, only genes expressed in at least two samples were retained to improve data quality (each group consists of 2 samples). DESeq2 was applied with sex as the experimental factor to normalize counts and model differential expression. Dispersion estimates were plotted to assess model fit. VST expression values were generated for downstream analysis. Expression data for selected lipid metabolism genes were then extracted for further analysis and visualization.

In both cases, selected genes involved in lipid metabolism were visualized using boxplots combined with overlayed beeswarm (beeswarm v0.4.0 package), which display individual sample variation colored by group. All analyses were conducted using R v4.4.0 and RStudio v2024.041+748.

## Statistical Analyses

All statistical analyses and graphing were performed using GraphPad Prism (v10). For comparisons between two groups, two-sided Student’s t-tests were used to determine statistical significance. For comparisons involving more than two groups, one-way ANOVA was employed. For comparisons between conditions in two or more groups, two-way ANOVA was used. P-values are reported in each figure, with a significance threshold of p < 0.05, where p > 0.05 is considered not significant (ns).

## Acknowledgments

We thank all members of the Goldberg lab for comments and intellectual discussion. The Goldberg lab is funded by R00AG058801 and DP2AI175641 to ELG, a pilot and feasibility award to ELG from the UCSF NORC P30DK098722, the Sandler Program for Breakthrough Biomedical Research, which is partially funded by the Sandler Program, and the Bakar Aging Research Institute (awards to both ELG and AL). The Benayoun lab is funded by the AthenaDAO student leader award (to CB) and NIA R01AG076433 (to BAB). The Stratman lab is funded by R35GM137976 awarded to ANS.

