## Supplemental Data for "Neutrophil TLR2 signaling promotes lipid accumulation and vascular plaque growth"

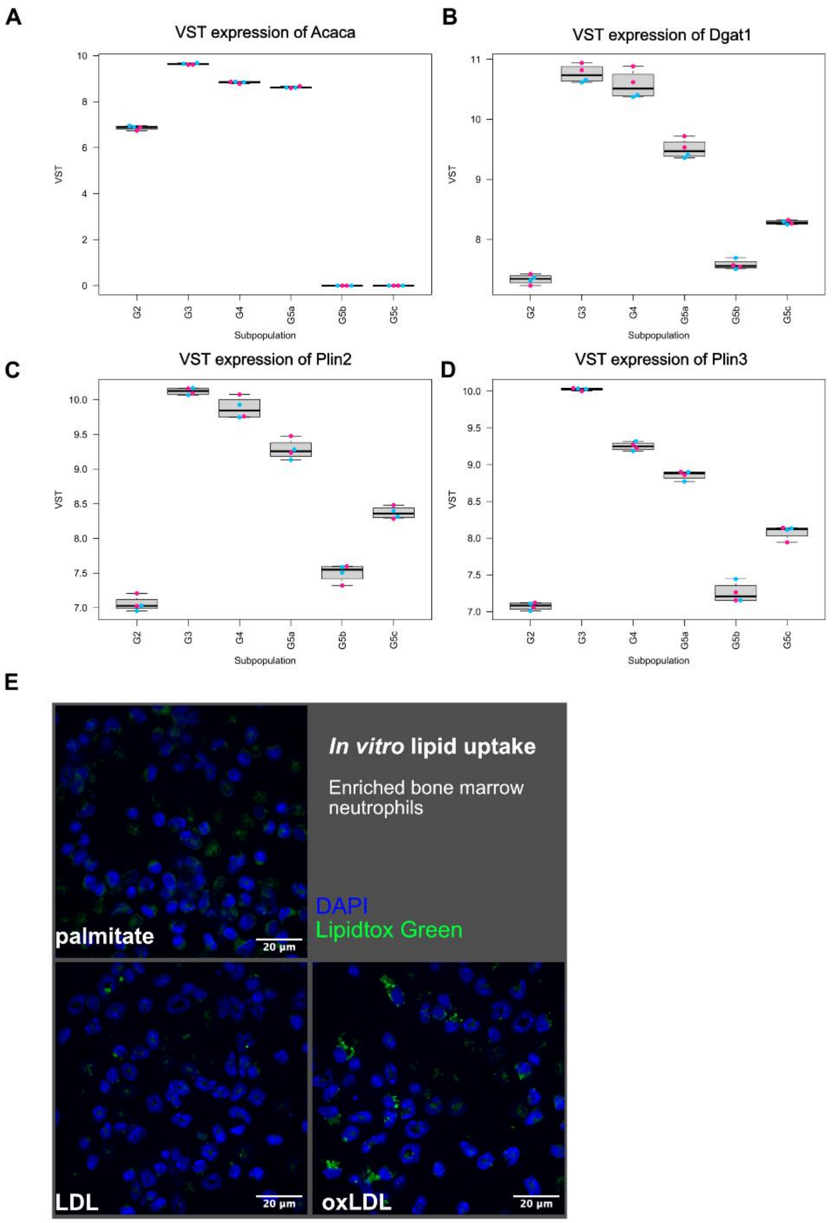


**Fig S1. Neutrophils are capable of lipid storage.**

(**A**) Single-cell RNA-seq data showing expression of fatty acid synthesis and lipid storage genes in across neutrophil maturation process. Pink represents female mice and blue represents male mice. (**B**) Confocal images of bone marrow enriched neutrophils treated overnight with the indicated non-fluorescent lipids and stained for Lipidtox Green and DAPI.

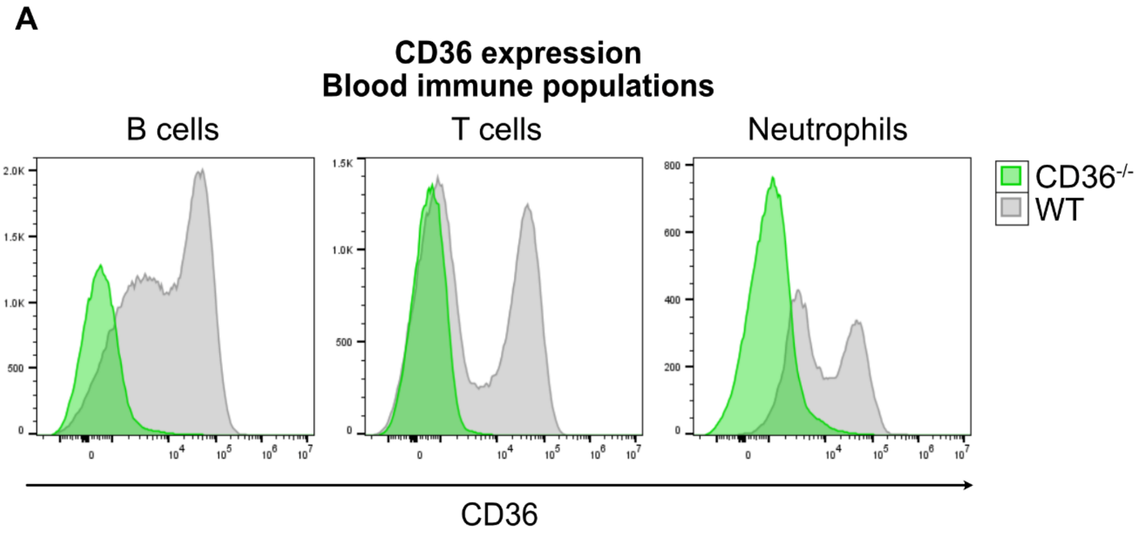


**Fig S2. Confirmation of CD36 deficiency in CD36-knockout mice.**

(**A**) Representative flow cytometry analysis of CD36 expression on B cells ($B220^+$), T cells ($CD3^+$) and neutrophils ($CD11b^+$ $Ly6G^+$) from the blood of WT and CD36 knock-out animals.

A

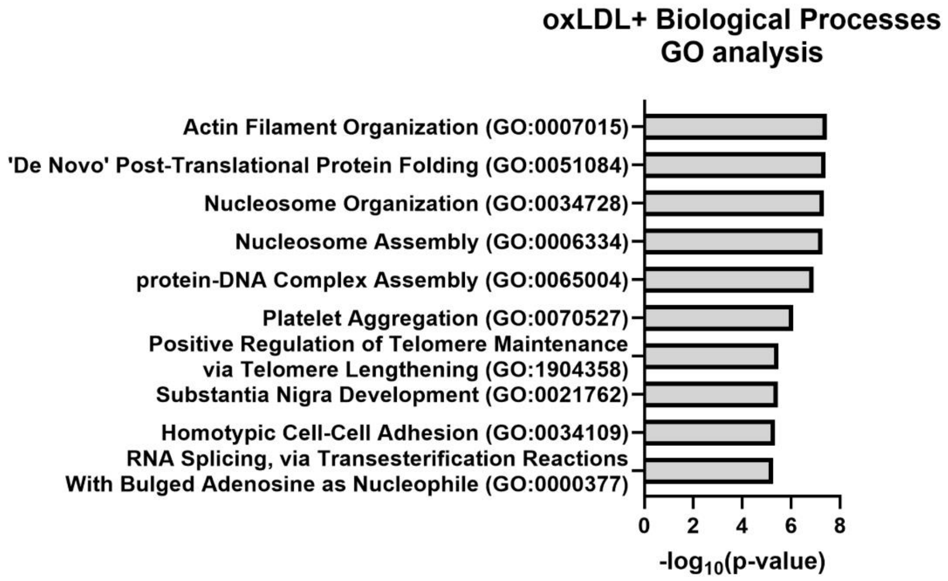


**Fig S3. Actin organization proteins are significantly different between oxLDL-positive and oxLDL-negative neutrophils.**

(**A**) Gene Ontology Biological Processes pathway enrichment analysis of proteins increased in $oxLDL^+$ neutrophils that were identified by mass spectrometry.

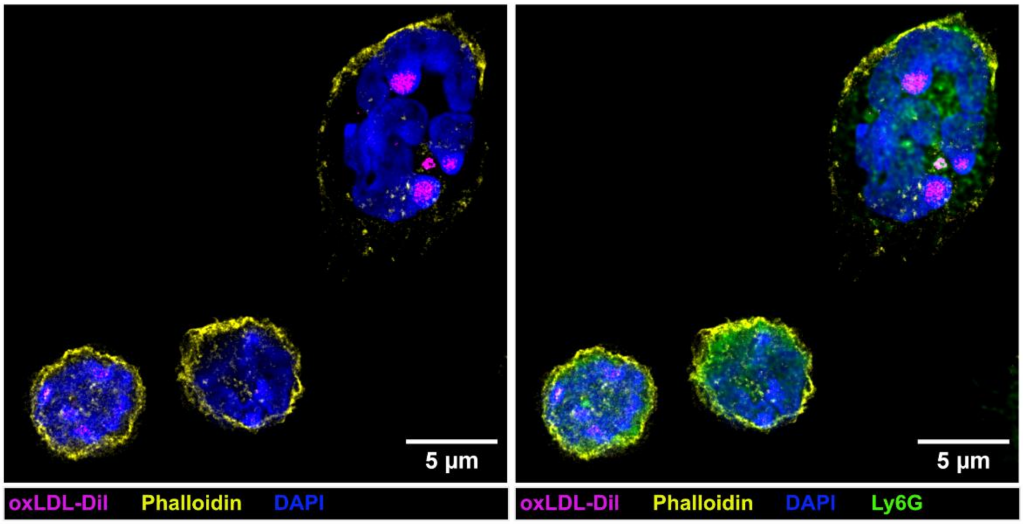


B

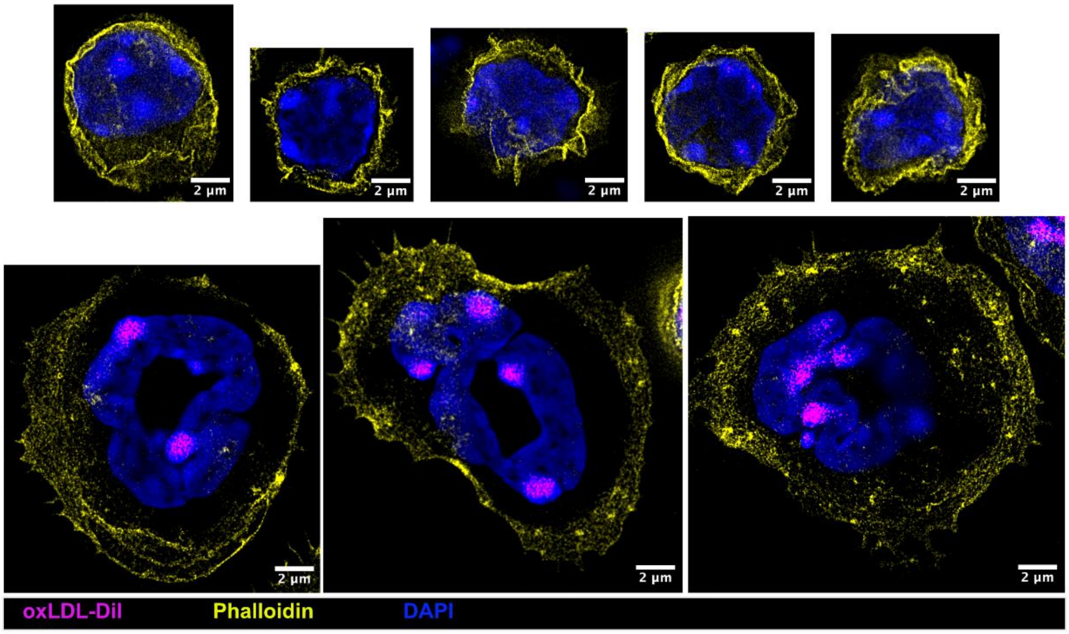


**Fig S4. F-actin polymerization is disrupted in neutrophils that have taken up oxLDL-DiI.**

(**A-B**) 100x STED imaging of zymosan-elicited peritoneal neutrophils that had high or low levels of oxLDL-DiI. (**A**) Multiple cells per field show selection of cells based on Ly6G stain for neutrophils (right panel). (**B**) Individual cells were imaged at super resolution to visualize actin cytoskeleton via Phalloidin stain. All cells were confirmed to have Ly6G stain (not shown for clarity). Phalloidin staining is shown in yellow, oxLDL-DiI is in magenta, and DAPI is shown in blue.

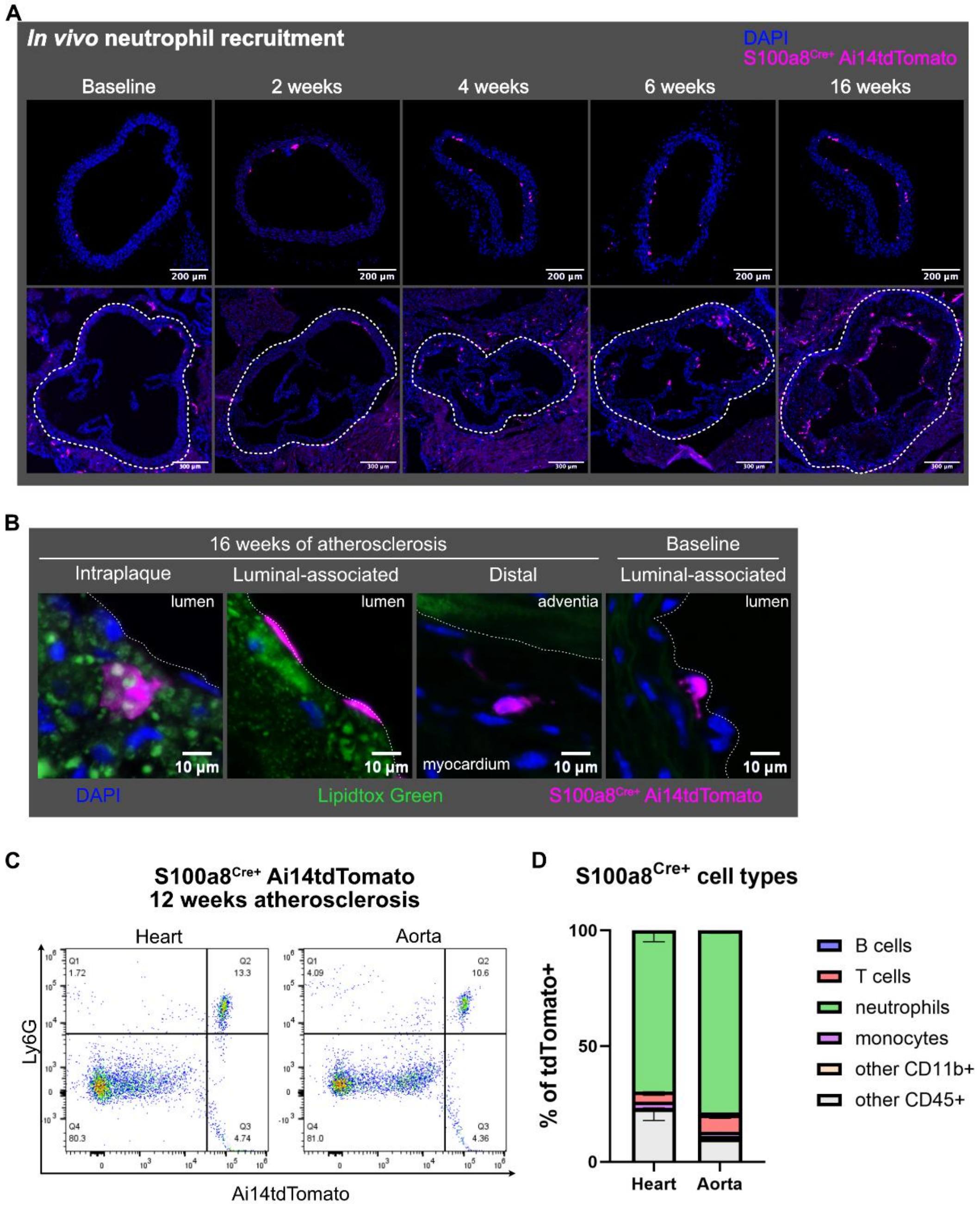


**Fig S5. Neutrophils accumulate in atherosclerotic lesions throughout disease.**

(**A**) Immunofluorescence images of the aorta cross-section (top) and the aortic root (bottom) at specified times after atherosclerosis induction in S100a8$^{Cre+}$ Ai14tdTomato mice. The aortic root region of interest is outlined with dashed lines. (**B**) Confocal images of 16-week and baseline aorta stained for Lipidtox Green. (**C**) Representative gating of CD45$^{+}$ cells and (**D**) quantified cell composition of S100a8$^{Cre+}$ Ai14tdTomato+ cells from heart and aorta at 12 weeks of atherosclerosis analyzed via flow cytometry.

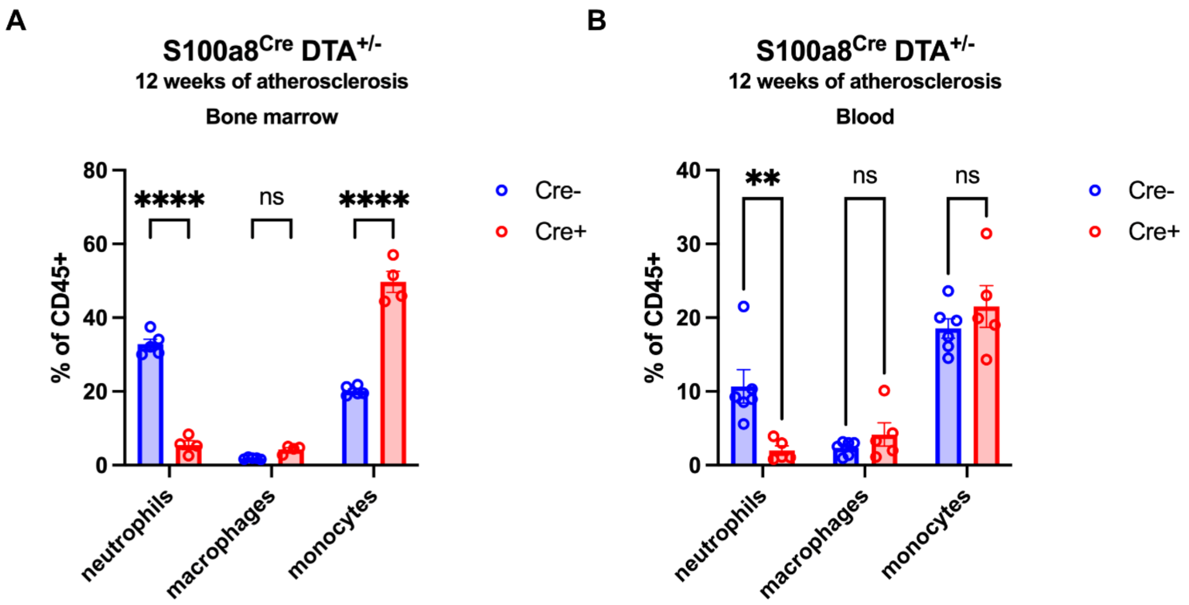


**Fig S6. Phagocyte characterization of atherosclerotic neutrophil-deficient mice.**

(**A**) Bone marrow and (**B**) blood immune characterization of S100a8$^{Cre+}$ DTA$^{fl/wt}$ and littermate control mice via flow cytometry. Cell populations were identified as neutrophils (CD11b$^{+}$ Ly6G$^{+}$), monocytes (CD11b$^{+}$, Ly6C$^{high}$, Ly6G$^{-}$) and macrophages (CD11b$^{+}$ F4/80$^{+}$). Data are from a representative mouse cohort from two independent experiments. Each symbol represents an individual animal. Statistical differences were calculated by two-way ANOVA. ** ≤ 0.001, **** ≤ 0.0001.

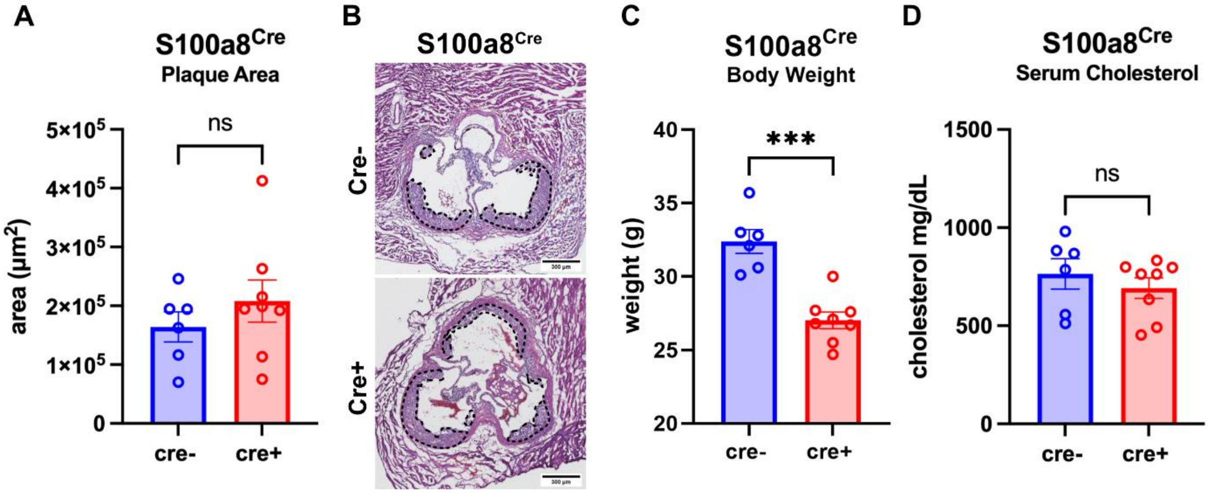


**Fig S7. Presence of the S100a8-Cre transgene does not influence plaque size.**

(**A**) Plaque area quantification, (**B**) representative images of H&E stain of aortic root, and (**C**) mouse body weights at 12 weeks of disease are shown for S100a8$^{Cre+}$ and littermate control mice. (**D**) Serum cholesterol quantification at 4 weeks of disease for the respective genotypes. Quantified plaques are highlighted with dashed lines. Data are from one representative mouse cohort, each data point on the graph represents a single animal. Statistical differences were calculated by t-test. *** $\leq$ 0.001
